# Isoform-specific translational control is evolutionarily conserved in primates

**DOI:** 10.1101/2023.04.21.537863

**Authors:** Jolene Draper, Julia Philipp, Zach Neeb, Richard Thomas, Solomon Katzman, Sofie Salama, David Haussler, Jeremy R. Sanford

## Abstract

Alternative splicing (AS) alters messenger RNA (mRNA) coding capacity, localization, stability, and translation. Here we use comparative transcriptomics to identify cis-acting elements coupling AS to translational control (AS-TC). We sequenced total cytosolic and polyribosome-associated mRNA from human, chimpanzee, and orangutan induced pluripotent stem cells (iPSCs), revealing thousands of transcripts with splicing differences between subcellular fractions. We found both conserved and species-specific polyribosome association patterns for orthologous splicing events. Intriguingly, alternative exons with similar polyribosome profiles between species have stronger sequence conservation than exons with lineage-specific ribosome association. These data suggest that sequence variation underlies differences in the polyribosome association. Accordingly, single nucleotide substitutions in luciferase reporters designed to model exons with divergent polyribosome profiles are sufficient to regulate translational efficiency. We used position specific weight matrixes to interpret exons with species-specific polyribosome association profiles, finding that polymorphic sites frequently alter recognition motifs for trans-acting RNA binding proteins. Together, our results show that AS can regulate translation by remodeling the cis-regulatory landscape of mRNA isoforms.

## Introduction

Comparative genomics and transcriptomics enable quantitative comparisons species and are powerful approaches for studying the evolution of gene regulatory mechanisms. For example, comparisons of tissue-specific gene expression and alternative splicing patterns across distantly related vertebrate species revealed that these modes of gene regulation evolve at different rates (Calarco et al. 2007; Mazin et al. 2018; Merkin et al. 2012; Barbosa-Morais et al. 2012). Comparative transcriptomic and proteomic analysis of primate cells also demonstrated that although steady-state protein levels are similar for orthologous genes, mRNA levels can vary dramatically. This suggests that overall mRNA levels might evolve under less rigorous evolutionary pressure compared to protein expression levels (Khan et al. 2013).

Isoform-specific mRNA expression may also be under selective pressure during evolution. Hundreds of regions in the human genome are ultraconserved with those of mouse and rat <u>(Bejerano et al. 2004)</u>. Many of these sequences overlap alternative exon sequences associated with nonsense mediated decay (NMD), suggesting that regulatory elements or the function of these mRNA isoforms are under strong purifying selection (Lewis et al. 2003; Pan et al. 2006; Lareau and Brenner 2015; Saltzman et al. 2008). Indeed, ablation or programmed mis-splicing of AS coupled NMD (AS-NMD) results in growth defects and loss of tumor suppression in cell culture and mouse embryo models (Thomas et al. 2020; Leclair et al. 2020). Taken together AS-NMD appears to be part of an evolutionarily conserved regulatory mechanism for fine tuning the expression of splicing factors.

Recently, several groups, including our own, discovered that mRNA isoforms can exhibit differential polyribosome association, suggesting that alternative splicing can be coupled to translational control (AS-TC) (Sterne-Weiler et al. 2013; Floor and Doudna 2016a; Blair et al. 2017; Reixachs-Solé et al. 2020; Wong et al. 2016). Although the mechanisms are poorly described, shuttling pre-mRNA splicing factors, such as Serine and Arginine-rich (SR) proteins and their exonic binding sites may play important roles in AS-TC (Sanford 2004; Maslon et al. 2014). The inclusion or exclusion of specific exons from mRNA isoforms may remodel the composition of cis-acting regulatory elements involved in translational control. Likewise, alternative 5’ and 3’ UTRs sequences are likely to harbor regulatory elements influencing isoform-specific translational control, such as upstream open reading frames (uORFs), internal ribosome entry sites (IRES), and RBP-binding sites (Pfeiffer et al. 2012; Gebauer et al. 2012). Thus, alternative splicing not only expands the protein coding capacity of eukaryotic genes but also influences the cytoplasmic fate of the resulting mRNA isoforms.

By contrast to AS-NMD, the physiological and evolutionary significance of AS-TC is unknown. In this study, we use comparative transcriptomics to test the extent of conservation of AS-TC and to identify functionally important exons that contribute to the coupling. We analyzed polyribosome associated mRNA isoforms from human, chimpanzee, and orangutan induced pluripotent stem cells (iPSCs) using subcellular <u>frac</u>tionation and high throughput <u>seq</u>uencing (Frac-seq). Among thousands of mRNA isoforms with differential polyribosome association, we discovered orthologous alternative splicing events subject to regulation by either primate-conserved with evidence for either conserved or species-specific translational control. We further showed that AS events with primate-conserved translational control show higher sequence conservation specifically in cassette exons, indicating the functional relevance of these exons. We validated the ability of orthologous exons from isoforms with differential polyribosome association to affect mRNA translation in vivo using luciferase reporters. Taken together our data suggest that exonic sequences can regulate isoform-specific mRNA translation and that AS-TC is a conserved mechanism of gene regulation across primates.

## Results

### Frac-Seq analysis of primate iPSCs

We previously developed Frac-seq as a method to determine the association of alternative mRNA isoforms with the polyribosome (Sterne-Weiler 2013). To test the hypothesis that alternative splicing coupled to translational control is conserved in primates, we applied the Frac-seq methodology to iPSCs of human, chimpanzee, and orangutan origins (Fig. 1A). We identified and quantified thousands of alternative splicing events from all subcellular fractions isolated from human, chimpanzee and orangutan iPSCs (Fig 1B). Of these events, about half of the alternative splicing events exhibited PSI values differing between fractions by more than 10%, indicating mRNA isoform-specific patterns of polyribosome association. We further separated these into events that generate non-productive NMD isoforms (AS-NMD) and events maintaining the integrity of their open reading frame. The latter we consider alternative splicing events that are potentially implicated in translational control (AS-TC events, Fig 1C). Of these, 35% exhibited differential sedimentation in both human and chimpanzee cells, whereas 12% of events were classified as AS-TC in all three species (Fig 1D).

**Fig 1.**
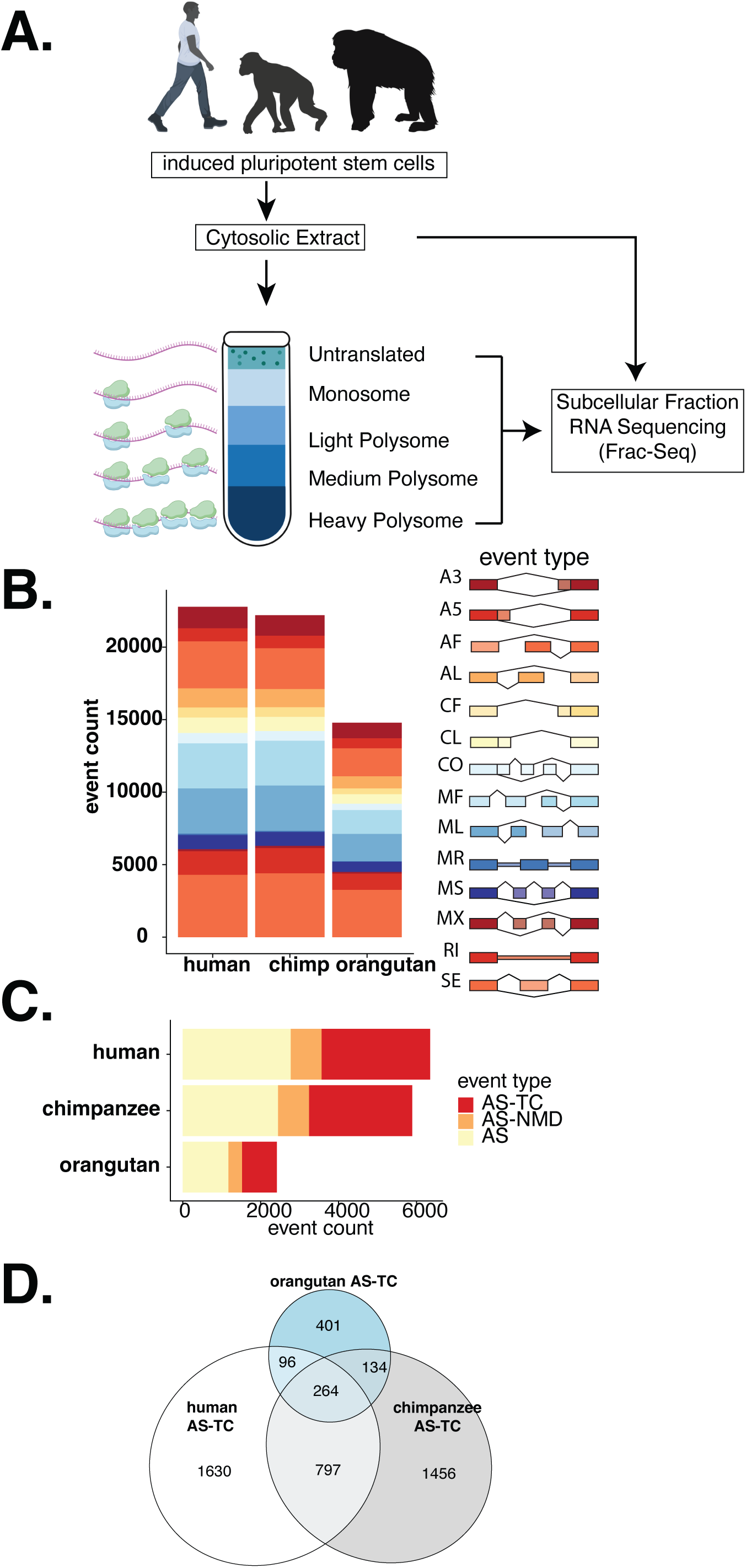
Frac-seq reveals polyribosome associated mRNA isoforms. A) Alternative splicing can influence multiple post-transcriptional regulation pathways. B) Frac-seq (subcellular fractionation and subsequent sequencing of polyA+ selected RNA from fractions) was performed on human, chimpanzee, and orangutan iPSCs. RNA from the total cytosolic lysate, the mono-ribosome (80s), the light (P2-P4), medium (P5-P8), and heavy polyribosome (P9+) was sequenced. B) Identification and quantification of alternatively spliced events was performed using junctionCounts. This pipeline allowed the identification of 14 different event types. C) Alternative splicing events were further classified into AS, AS-TC, and AS-NMD events. The proportions of these three event groups are comparable between the three cell lines. D) Out of the events categorized as either AS, AS-TC, or AS-NMD, about 900 events were identified in all three cell lines. E) Out of the events categorized as AS-TC, over 300 events were identified in all three cell lines.

### AS-TC is conserved across primate evolution

To identify patterns of polyribosome association in orthologous AS-TC events, we performed unsupervised hierarchical clustering on the Spearman correlation of the mean PSI for each species, across all fractions. As a control, we performed the same analysis using orthologous splicing events exhibiting uniform polyribosome association (ΔPSI <10%). These events cluster predominantly by species rather than by subcellular fraction (Fig 2A). By contrast, orthologous AS-TC events show a stronger correlation for PSI between subcellular fractions rather than by species (Fig 2B), indicating that isoform-specific sedimentation profiles are conserved between species. To identify orthologous isoforms with the most and least similar sedimentation profiles we calculated the difference in PSI between pairs of species and summed those differences into a single distance metric. The Manhattan distance for all orthologous AS-TC events results in a right-skewed distribution where the far left and right tails represent the AS-TC events with the most and least conserved sedimentation profile, respectively (Supp Fig 1A). We visualized these subsets of events in correlation heatmaps. The colors indicate the Spearman correlation of PSI values in events with low cumulative distance (similar sedimentation profiles) (Fig 2C) and in events with high cumulative distance (distinct sedimentation profiles) (Fig 2D).

**Fig 2.**
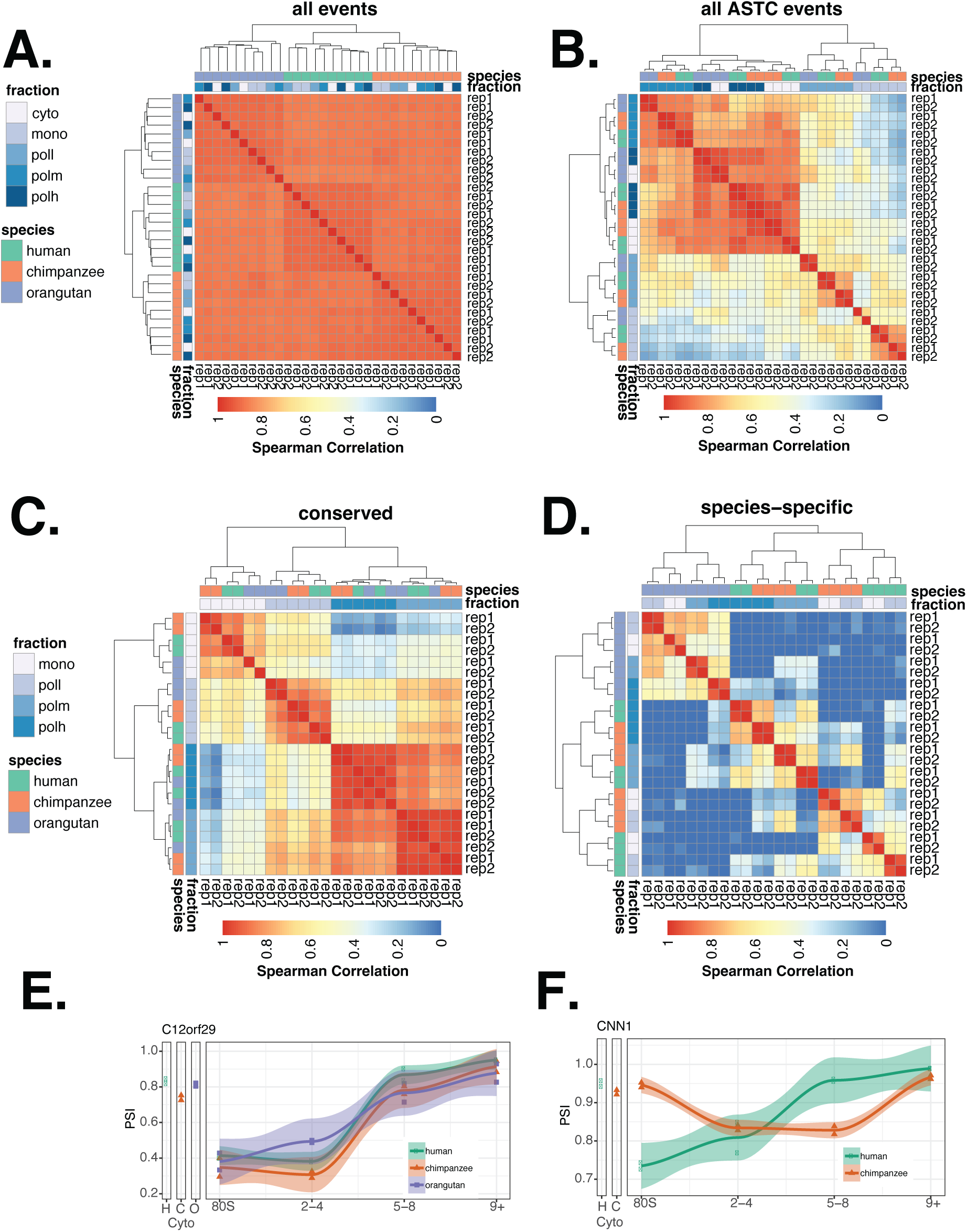
Orthologous AS-TC events exhibit either conserved or species-specific sedimentation profiles. A) Heatmap of Spearman correlation of PSI values of all identified orthologous events. The columns and rows represent the total cytosolic lysate and the 4 subcellular fractions in each cell line. The colors represent the Spearman correlation of PSI values between pairs of fractions (red = high correlation, blue = low correlation). B) Heatmap of Spearman correlation of PSI values of all orthologous AS-TC events. C,D) Heatmaps of Spearman correlation of PSI values of orthologous AS-TC events. C) The events in this heatmap exhibit sedimentation profiles consistent across all three cell lines as shown in the example in panel E. D) The events in this heatmap exhibit species-specific sedimentation profiles as shown in the example in panel F. E) The skipped exon event within C12orf29 is an example of an alternative splicing event with conserved sedimentation profiles across all three cell lines. F) The alternative first exon event of CNN1 is an example of an alternative splicing event with species-specific sedimentation profiles.

Interestingly, events with similar sedimentation profiles cluster by fraction, indicating the PSI values in these fractions are more similar to each other than to the other fractions within the same species. By contrast, in events with distinct sedimentation profiles, the different polyribosome fractions from each species cluster together, indicating higher similarity within the species than across the fractions. For example, C12orf29 exhibits a similar sedimentation profile between species (Fig 2E, left panel), whereas CNN1 alternative splicing generates isoforms with species-specific sedimentation patterns (Fig 2D, right panel). In events with species-specific sedimentation patterns, the PSI values across the polyribosomal fractions differ between species, despite a well correlated nuclear output, represented by the PSI value in the cytoplasmic fraction (r= 0.90, 0.85, and 0.86 for human vs. chimp, human vs. orangutan and chimp vs orangutan, respectively. Supplemental Fig 2).

### Exon sequences associated with AS-TC are strongly conserved

To test the hypothesis that exonic sequence elements are determinants of AS-TC, we examined the sequence conservation of orthologous skipped exons (SE) mRNA isoforms with similar or species-specific polyribosome profiles. We also visualized the phastCons score for AS events that were not associated with differential polyribosome association. As expected, skipped exons that are not associated with AS-TC are much less conserved compared to their flanking exons. By contrast, exons linked to AS-TC exhibit significantly higher phastCons scores and are more similar to their flanking exons. Further, AS-TC exons that have conserved sedimentation profiles between species have elevated phastCons scores relative to the other classes (Fig 3A). The high degree of sequence conservation within AS-TC exons suggests the presence of functional elements within these exons that influence polyribosome association. We next investigated the relationship between sequence variation and isoform-specific polyribosome sedimentation. Using pairwise alignments, we compared human and chimpanzee AS-TC cassette exons and calculated the SNP frequency within AS, conserved AS-TC and species-specific AS-TC events. After normalizing for sequence length, we observe the highest single nucleotide variants (SNV) frequency in AS-TC exons with species-specific polysome association patterns, followed by canonical AS exons, and then by AS-TC exons with conserved sedimentation (Figure 3B). As expected, SNVs are enriched of variants around 50bp away from the splice sites in the exon sequence in SE events (Figure 3 C,D). This enrichment agrees with previously observed positional biases of SNPs around splice sites (Majewski and Ott, 2002). Interestingly, we observed a lower SNP density in AS-TC exons with similar sedimentation profiles between species. Taken together, these data suggest that exonic sequence variation can influence polyribosome association of mRNA isoforms.

**Fig 3.**
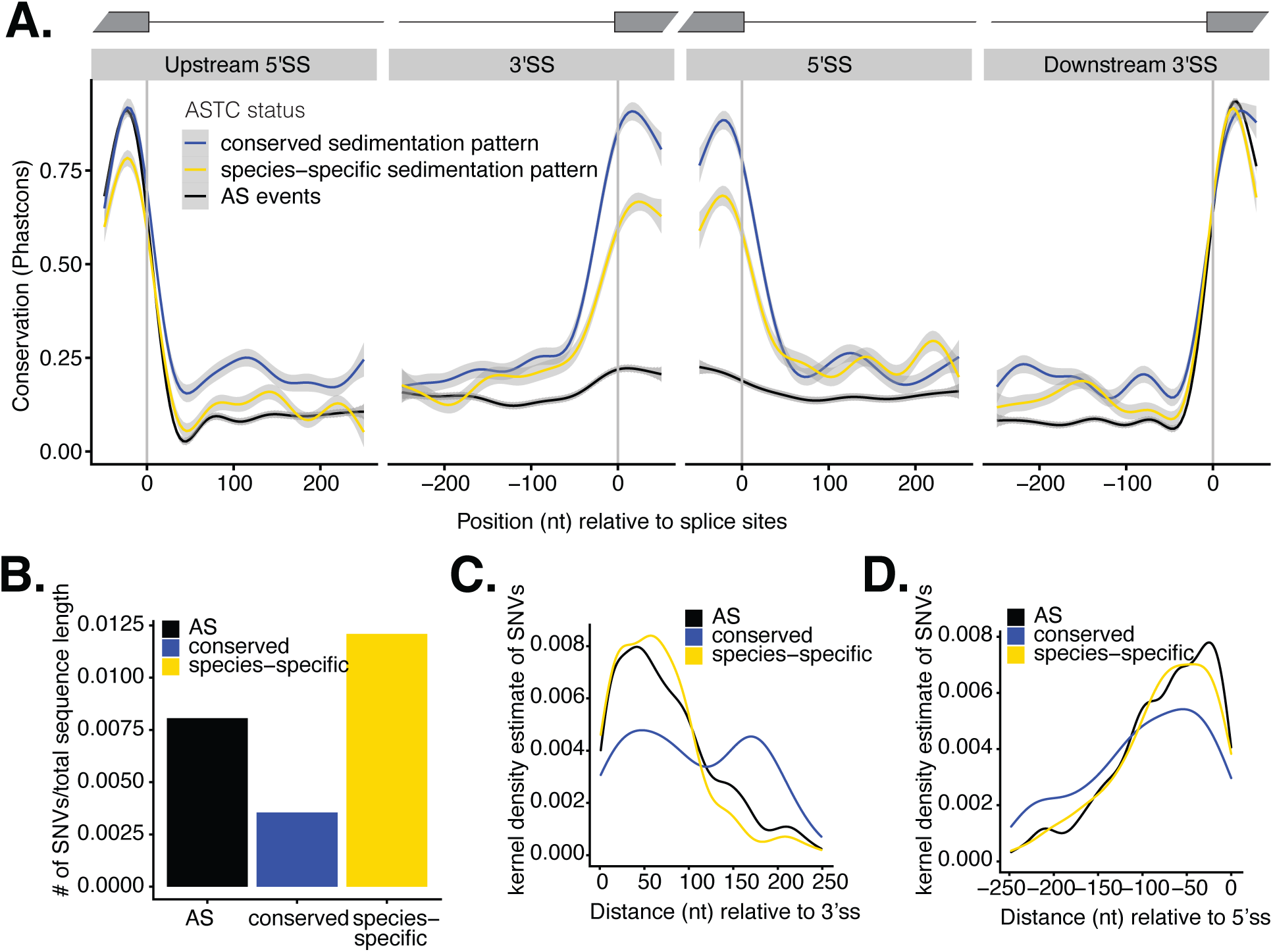
Alternative splicing events with sedimentation profiles consistent across species show higher sequence conservation. A) Sequence conservation of exon/intron boundaries of AS-TC and AS skipped exon events represented by phastCons scores. A lower score indicates less conservation. AS-TC events with conserved sedimentation profiles in blue, AS-TC events with species-specific sedimentation profiles in yellow, and simple AS events in black. B) Length normalized SNV density for skipped exon sequences associated with AS (black), conserved AS-TC profiles (blue) or species-specific AS-TC profiles (yellow). C,D) Positional distribution of SNP density in skipped exon sequences associated with AS (black), conserved AS-TC profiles (blue) or species-specific AS-TC profiles (yellow) relative to the 3’ or 5’ splice site, respectively.

### Single nucleotide variants in orthologous AS-TC exons influence translation

To test the hypothesis that sequence differences associated with isoform-specific sedimentation patterns regulate mRNA translation, we created translational luciferase reporters from SE events from exhibiting AS-TC. Figure 4A shows the schematic for the skipped exon luciferase reporters, where either the human (green) or chimpanzee (orange) skipped exon were inserted in-frame, upstream of the firefly luciferase. For each reporter, we measured resulting luciferase activity and steady-state mRNA levels. Frac-seq analysis revealed differential sedimentation profiles for human and chimp isoforms from the GGCX, MELK and SUMF2 genes (Fig 4C-E left panel, respectively). These results were particularly striking as the orthologous exon sequences for each of these events differ by only a single nucleotide (Supplemental Figure 3). Dual luciferase assays revealed that orthologous reporters for GGCX and MELK reporters exhibited significantly different activity (Figure 4C-D, middle panels). Frac-seq data indicates that human GGCX exon 2 inclusion is enriched in polyribosome fractions compared to chimp, where it is broadly distributed across the gradient. Consistent with this result, the human GGCX luciferase reporter is expressed more robustly than the chimp reporter. The opposite trend is observed for MELK exon 3, which is depleted from polyribosome fractions relative to the orthologous chimp isoform. In this case, the luciferase reporter containing the chimp sequence is more robustly expressed than the orthologous human reporter (Figure 4C-D, middle panels). This effect was likely due to translational control, as the steady-state mRNA levels are do not differ significantly between the orthologous GGCX and MELK reporters (Figure 4 C-D, right panels). Additionally, the stability of luciferase enzyme levels following translational arrest with the translation elongation inhibitor cycloheximide are not significantly different for the orthologous reporters, indicating that the single nucleotide substitution does not impact protein stability (supplemental figure 4). By contrast, orthologous reporters from SUMF2, exhibit nearly identical luciferase expression and reporter mRNA levels. These data are consistent with the Frac-seq profiles for human and chimp SUMF2 isoforms, which exhibit increased exon 2 inclusion in both species. Taken together, these data demonstrate that single nucleotide polymorphisms can lead to large changes in translational control.

**Fig 4.**
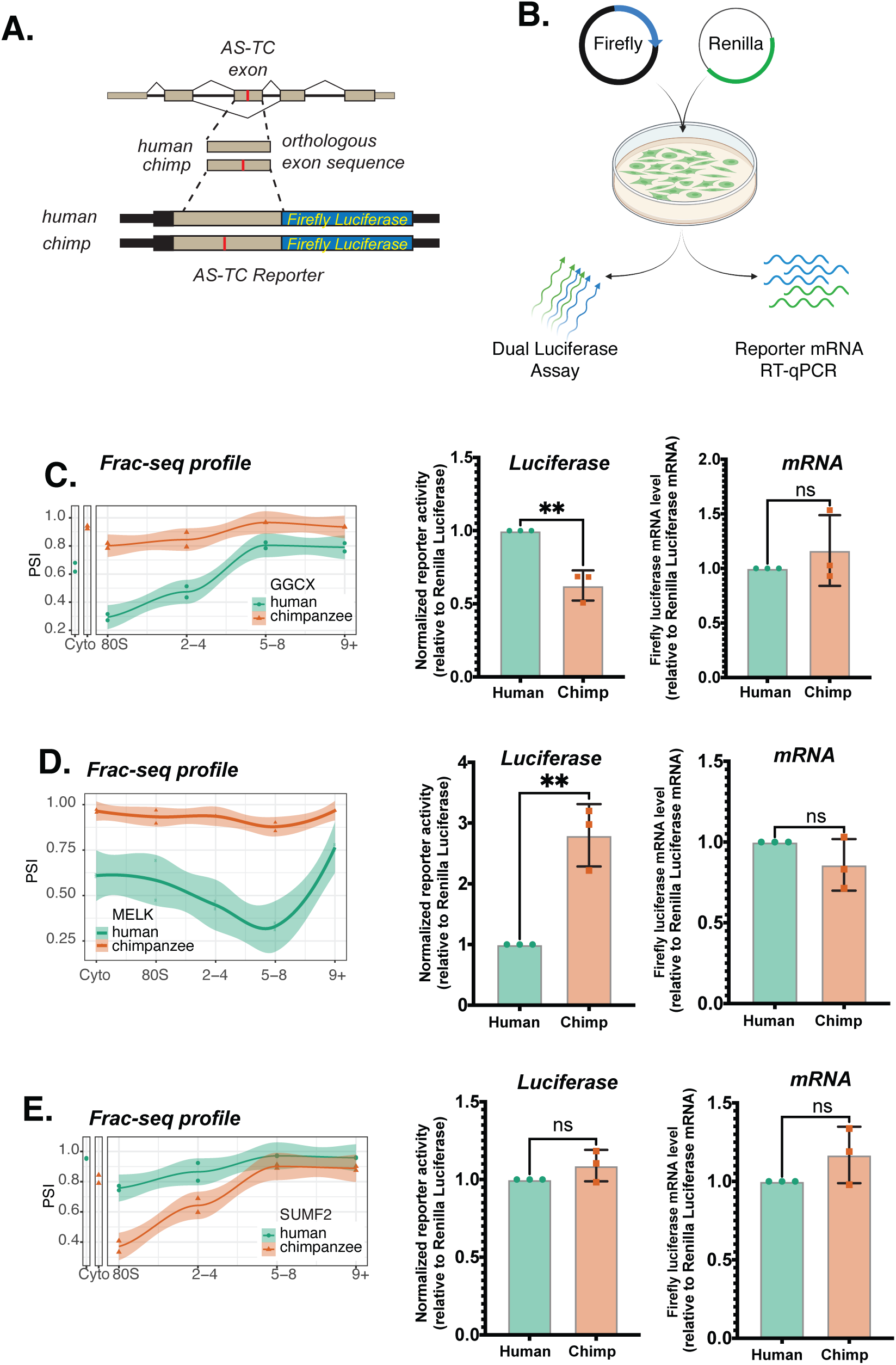
AS-TC cassette exons drive isoform-specific expression. A) Schematic diagram of the pairs of luciferase reporter constructs containing either the human (green) or chimpanzee (orange) cassette exon from different genes exhibiting AS-TC. B) Schematic diagram of dual luciferase assay and RT-qPCR assay. C-E) Polyribosome sedimentation profiles from human and chimpanzee iPSCs (left panel), dual luciferase reporter assay data (middle panel) and RT-qPCR data (right panel) for isoforms from the GGCX, MELK and SUMF2 genes, respectively. Dual luciferase reporter assay and RT-qPCR analyses were performed on three independent transfections of orthologous reporter pairs. Statistical analysis by one-way ANOVA.

### Alternative splicing alters the cis-regulatory landscape of mRNA isoforms

In order to test the effect of alternative splicing on the cis-regulatory landscape of mRNA isoforms, we performed pairwise alignments of alternatively spliced exons (e.g. skipped exons) between two species at a time (human-chimpanzee and human-orangutan) and identified the sequence differences, predominantly SNVs, between the species. We then tested the effect of these sequence differences on predicted protein-RNA interactions sites. For that purpose, we scored the similarity of RNAcompete-derived positional weight matrices (PWMs) (Ray et al. 2017) against the sequences surrounding SNVs in a sliding window and collected a predicted binding score for each tested RBP (Figure 5A). We performed this analysis on three groups of alternatively spliced events: (i) events that are classified as AS-TC in one species and only AS in the other, (ii) events that are AS-TC in both species, and (iii) AS-only events. Next, we aimed to identify changes in the cis-regulatory landscape of mRNA isoforms that might be connected to differential, conserved polysome association or species-specific polysome association. For that, we compared the frequency of RBP binding sites that are changed in binding score due to the presence of a SNV in the different event groups (Figure 5B, C), whereas the AS events in both species serve as a control. Interestingly, both human-chimpanzee and human-orangutan sequence differences resulted in changes in putative RBP binding sites with frequencies significantly different to the AS-only control group (Figure 5B,C). Out of the 10 PWMs that showed significant differences in frequency between the human-chimpanzee group and the 21 in the human-orangutan group, two were present in both analyses (Figure 5D). The AS events analyzed for the human-chimpanzee and the human-orangutan comparison are predominantly distinct sets, with an overlap of 205 events (Figure 5E). Taken together, these data suggest that SNPs associated with AS-TC exons with divergent Frac-seq profiles may alter the binding sites for RBPs associated with translational control.

**Fig 5.**
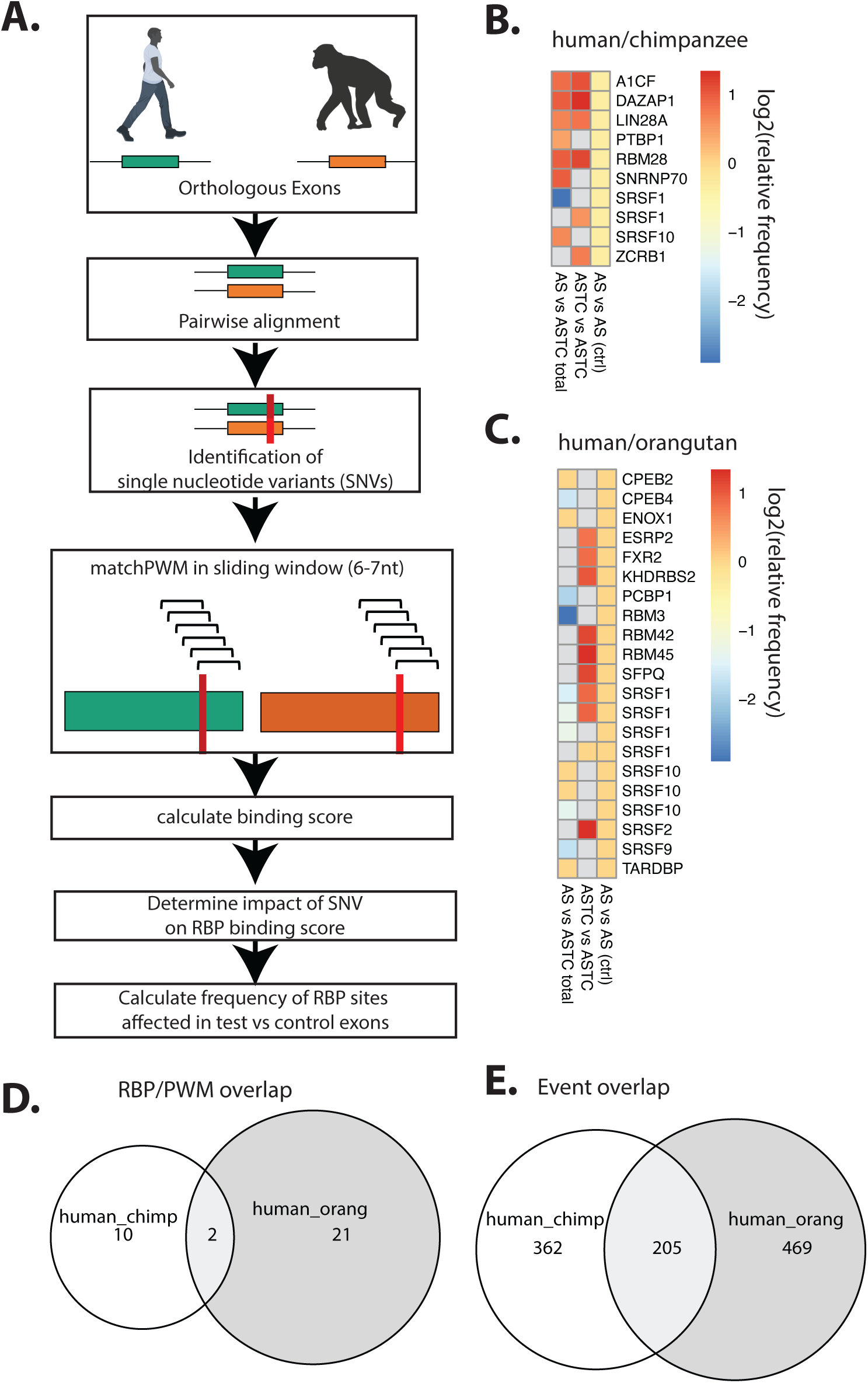
Alternative splicing alters the cis-regulatory landscape of mRNA isoforms. A) Schematic diagram of RBP binding analysis. B) Heatmap of log2 of relative frequencies of predicted RBP binding sites changed through single nucleotide variants between human and chimpanzee skipped exon sequences in AS/AS-TC events, AS-TC/AS-TC events and AS/AS events compared to the AS/AS event control group. C) Heatmap of log2 of relative frequencies of predicted RBP binding sites changed through single nucleotide variants between human and orangutan skipped exon sequences in AS/AS-TC events, AS-TC/AS-TC events and AS/AS events compared to the AS/AS event control group. D) Euler diagram depicting the overlap of RBPs with significant differences in frequencies compared to the background between the human/chimpanzee and the human/orangutan analysis. E) Euler diagram depicting the overlap of analyzed events in the human/chimpanzee and the human/orangutan analysis.

## Discussion

Here, we applied a comparative transcriptomics approach to determine how genetic variation influences mRNA isoform polyribosome association in closely related primate stem cell models. We hypothesized that divergent sequences may change the cellular fate of orthologous mRNA isoforms and unveil functional elements important for post-transcriptional regulation of gene expression. Using Frac-Seq, we identified orthologous mRNA isoforms with similar or distinct distributions in sucrose gradients. At face value, these data demonstrated that polyribosome association can be regulated in an isoform-specific manner across primates. Using exon skipping events as a model, we found that orthologous isoforms with similar polyribosome sedimentation profiles exhibited strong sequence conservation across vertebrate phylogeny when compared to genes with species specific sedimentation profiles or non-AS-TC skipping events. We present evidence that single nucleotide polymorphisms within skipped exons can be sufficient to alter reporter mRNA translation. These data point to the existence of sequence elements within coding exons that contain the capacity to regulate translation. We tested this hypothesis by leveraging publicly available RBP recognition motifs to investigate the impact of the SNVs on potential protein-RNA interactions (Ray et al. 2013). This analysis revealed several *trans*-acting RBPs that recognize sequences associated with AS-TC, suggesting that mRNP composition influences the cytoplasmic fate of mRNA isoforms.

Our data show that the coupling of alternative splicing to translational control is conserved across multiple species of primate cell lines, representing approximately 13 million years of primate evolution. We discovered 1,000-3,000 AS-TC events in human, chimp, and orangutan iPSC lines. Approximately 30-50% of AS-TC events were classified as AS-TC in another primate cell model and 12% alternative events exhibited AS-TC in all three species (Figure 1E). By comparing PSI between fractions and species, we found that isoform abundance was strongly correlated by fraction, reflecting similar isoform-specific sedimentation patterns between species (Figure 2B). AS-TC events with shared sedimentation profiles also showed stronger alternative sequence conservation compared to species-specific AS-TC or AS events (Figure 3). The evolutionary conservation of these sequences indicates a potentially important biological function for gene regulation by AS-TC and/or points to the presence of potentially important *cis*-regulatory elements.

The mechanisms coupling alternative splicing to translational control are unclear. Whereas much is known about how alternative promoter usage and alternative polyadenylation signal sites can regulate isoform-specific translation (Niederer et al. 2022; Floor and Doudna 2016b; Blair et al. 2017)., the ways in which alternative splicing of coding exons influences translational control are less clear. By analyzing orthologous skipped exons with similar or divergent sedimentation profiles in sucrose gradients, we found significant differences in nucleotide sequence conservation. Indeed, orthologous alternative exons with similar sedimentation profiles between species exhibited a high degree of sequence conservation compared to exons with divergent sedimentation patterns (Fig. 3). Using a previously described luciferase reporter system, we found that a single nucleotide substitution induced differences in reporter mRNA translation in HEK 293T cells (Fig. 4). For example, GGCX exon 2 and MELK exon 3 show different patterns of exon inclusion across sucrose gradients in human and chimp iPSCs. These differences in Frac-seq profiles are reflected in allele-specific expression of luciferase reporters *in vivo*. Together, these data argue that cis-acting elements within the open reading frame provide another regulatory point for translational control on the level of alternative splicing.

How can a single nucleotide change in an exon impact the translational control message it resides in? One possibility is that the reach of exonic splicing enhancers extends to the cytosol. We previously demonstrated that the nucleo-cytoplasmic shuttling SR protein SRSF1 and a well characterized recognition motif from the *FN1* pre-mRNA can enhance translation both *in vitro* and *in vivo* (Sanford et al. 2004). SRSF1 promotes translation initiation through interactions with mTORC1 components and inducing phosphorylation of the translational repressor EIF4E-BP (Blaustein et al. 2005; Michlewski et al. 2008); (Sanford 2004; Maslon et al. 2014)). Based on evidence presented here, we hypothesize that the sequences of AS-TC exons interact with trans-acting RNA binding proteins to drive isoform-specific translational control. We found that recognition motifs for several splicing factors such as SRSF1 and SRSF10 as well as translational control factors including LIN28A, FXR2, and DAZAP1 are frequently altered by SNPs within orthologous AS-TC exons with divergent sedimentation profiles (Figure 5). Future work to determine the precise mechanisms as to how exons such as MELK exon 3 or GGX exon 2 regulate translation will be needed to understand how coding sequences establish translational control.

Several mechanisms were not explored in our analysis and remain viable options for specific transcripts. For example, changes in mRNA sequence could contribute to isoform-specific differences in RNA secondary structure, which has previously been shown to regulate mRNA translation (Mao et al. 2014; Chemla et al. 2020; Mauger et al. 2019). For example, more stable secondary structures near the start codon attenuate translation initiation (Kudla et al. 2009). Likewise, codon optimality influences translation elongation and RNA stability in a variety of species (Gu et al. 2010; Tuller et al. 2010; Presnyak et al. 2015). In metazoans, codon optimality is important for enhancing the expression of intronless genes and can also influence codon usage within alternatively spliced isoforms by various mechanisms, including inducing frameshifts, modulating GC content, or by varied use of exons with suboptimal codons. Despite these open questions, the work presented here indicates that regulation of gene expression by AS-TC is evolutionarily conserved across the great apes and implicates RNA binding proteins as potential regulators of AS-TC in primates. Taken together, these findings suggest that alternative splicing influences mRNP composition by creating complexes with isoform-specific combinations of RNA binding proteins.

## Materials & Methods

### iPSC generation and culture

Human iPSC line C305 was derived from the GM12878 lymphoblastoid cell line using a proprietary episomal, plasmid-based reprogramming method by Cellular Dynamics. C305 cells were cultured on matrigel in mTeSR. Integration-free chimpanzee (Eip-8919-1A) and orangutan (Jos-C31) iPSCs were generated from primary fibroblasts as previously published by Field et al. (Field et al. 2017) and cultured on vitronectin in Essential 8 media.

### Fractionation, polyribosome profiling, RNAseq

Frac-seq experiments were performed as previously published (Sterne-Weiler et al. 2013) using human, chimpanzee, and orangutan iPSC lines, with modification to the number of fractions collected. Cytosolic extracts from cell lines/tissues are fractionated by sucrose gradient centrifugation. We collected the total cytosolic lysate, the monoribosomal fraction (80s), as well as light (P2-4), medium (P5-8), and heavy (P9+) polyribosomal fractions. We then polyA+ selected RNA from each fraction and individually barcoded libraries using the Bioo NEXTflex Directional RNASeq kit. Multiplex-pooled libraries were sequenced on an Illumina HiSeq 4000 using paired-end 2×150bp sequencing, resulting in 75-150M reads per sample (Supplementary Figure 1B) with approximately 40-50% junction reads per sample.

### Mapping of Illumina short read RNA sequencing

The reads were mapped to the human genome assembly hg38, the chimpanzee (Pan troglodytes) genome assembly panTro6, and the sumatran orangutan (Pongo abelii) genome assembly ponAbe3 using STAR v2.7 (Dobin et al. 2013). Repeat sequences were masked by mapping to repeatMasker sequences (Smit et al. RepeatMasker Open-4.0 at http://repeatmasker.org) using Bowtie2 (Langmead et al. 2009). PCR duplicate removal was performed by collapsing fragments with common start and end positions and CIGAR strings using in house scripts. All data collection and parsing was done with bash and python2.7. Statistical analyses and data visualization were performed using R programming language version 3.5.1.

### Identification and quantification of orthologous alternative splicing events

We used Stringtie (Pertea et al. 2015) to identify unannotated transcripts and used CAT transcriptomes (Fiddes et al. 2018) (for human (hg38), chimpanzee (panTro6), and orangutan (ponAbe3)), together with the stringtie merge command to generate the final transcriptomes for each species. Pairwise alternative splicing events were identified by pairwise comparison of all transcripts with at least one exon-intron junction in common. An alternative event was defined to be a set of exons unique to one transcript that are surrounded by two exon-intron junctions common to both transcripts or by one junction and the transcript terminus.

Alternative splicing events were quantified by counting the number of reads supporting the exon-exon junction and the number of reads supporting the exon-intron junction of each event. PSI (percent spliced in) values were calculated as the ratio of number of reads supporting the included isoform to the number of reads supporting both isoforms.

The event identification and quantification was implemented using in house python scripts. Orthologous alternative splicing events were identified by mapping the event sequences from one species to the genomic sequences of all others. The genes containing the orthologous event coordinates were determined using the CAT transcriptome annotations.

### Cross-fraction and Cross-species comparison of conserved and species-specific orthologous events

All events identified were filtered to be supported by at least 15 junction reads (per comparison). Alternative splicing events undergoing translational control, or AS-TC events, were defined as events with a change in PSI value (delta PSI) between any two adjacent fractions of at least 0.1. Consequently, alternative splicing events not undergoing translational control were defined as events with a minimum PSI > 0 and delta PSI < 0.1. Alternative splicing events leading to nonsense-mediated decay (AS-NMD) were identified using *in silico* translation of raw transcripts and subsequent identification of premature termination codons (PTCs) (technically CDSinsertion).

For estimating the difference/conservation of the polysome association pattern we calculated the Manhattan distances for each event between each two[1] [2] species. The Manhattan Distance is the sum of differences in mean psi between two species across all fractions. Min/max normalization of the Manhattan distance allowed us to identify events with overall different sedimentation profiles as opposed to events with similar sedimentation profiles at a different y-axis intercept. We ranked all AS-TC and AS-NMD events based on their min/max normalized Manhattan distance and used the top and bottom 10% (= 350 events) for further analysis, considering them the least and most conserved set of events respectively.

### Determination of Sequence conservation

To determine the sequence conservation of AS and AS-TC events with conserved or species-specific sedimentation profiles, phastCons (Siepel and Haussler; Siepel 2005) scores were obtained from the UCSC genome browser (Kent et al. 2002). For skipped exon events, phastCons scores were obtained for 100nt windows around both splice sites of the cassette exon as well as around the upstream 5’ss and the downstream 3’ss. [3] [4] The scores were visualized with local nonlinear smoothing using a generalized additive model.

### RNA purification and RT-qPCR

Total RNA was isolated using the Direct-zol RNA MiniPrep Kit (Zymo Research). 800ng of the RNA were treated with RQ DNase (per protocol). The DNase (Promega) treated RNA was reverse transcribed using the High-Capacity cDNA reverse transcriptase kit (Applied Biosystems). 1:200 dilutions of the cDNA were made. For the qPCR, we used Luna 2x SYBR premix (total volume 20*μ*l per reaction), 0.25 nM primers and 5*μ*l diluted cDNA. qPCR was performed on QuantStudio 3 Real-Time PCR System (Applied Biosystems, Thermo Fisher) according to MIQE guidelines (Bustin et al. 2009).

### Luciferase Reporters

Luciferase activity was assayed 24 hours post transfection using Dual-Glo Luciferase Assay System (Promega). For a 6 well plate, transfections were performed with lipofectamine with either 2*μ*g pLCS plasmid (previously published Sanford et al.) plus 125ng control plasmid (rluc) (for skipped exon events) or 1*μ*g p5UTR (pLightSwith_5UTR, from Switch Gear) 1*μ*g plus 250ng control plasmid (pmir) (for alternative first exon events) per well.

### Alignment and identification of SNVs and indels

The sequences of alternative regions of AS-TC events from human and chimpanzee were globally (Needleman-Wunsch alignment) aligned using R Biostrings (Pagès et al. 2020) (function pairwiseAlignment) using default settings. Mismatches, insertions, and deletions were identified in the pairwise alignments using R Biostrings.

### RBP binding prediction

RNAcompete (Ray et al. 2009, 2013) data sets available on ENCODE (ENCODE Project Consortium 2012) were used to predict RBP binding sites affected by single nucleotide sequence differences between human and chimpanzee AS-TC events. The matchPWM (Wasserman and Sandelin 2004) function from the R Biostrings package (Pagès et al. 2020) was used to score PWMs based on the RNAcompete data in a sliding window across the identified sequence differences. Matches achieving at least 80% of the maximum score were recorded for both human and chimpanzee datasets. The matches were compared between the species in form of a deltaPWMscore (e.g. PWMscore(human) - PWMscore(chimp)) significance was determined using Chisquare test for proportions.

## Data Access

All raw and processed sequencing data generated in this study have been submitted to the NCBI Gene Expression Omnibus (GEO; https://www.ncbi.nlm.nih.gov/geo/) under accession number GSE230441.

## Acknowledgements

We would like to thank our colleague Dr. Gina Mwala for critical feedback on the manuscript and Profs. Joshua Arribere, Angela Brooks, and Benedict Patten and Dr. Tim Sterne-Weiler for thoughtful discussion. We wish to thank Erin LaMontagne, and Lila Whitehead for their help culturing primate iPSCs. This research was made possible by a grant from the California Institute of Regenerative Medicine (GCIR-06673-A to JRS) and from the National Institutes of Health (R35GM130361 to JRS). The contents of this publication are solely the responsibility of the authors and not necessarily the official views of CIRM or any other agency of the State of California.

---

**Supplementary Fig 1.**
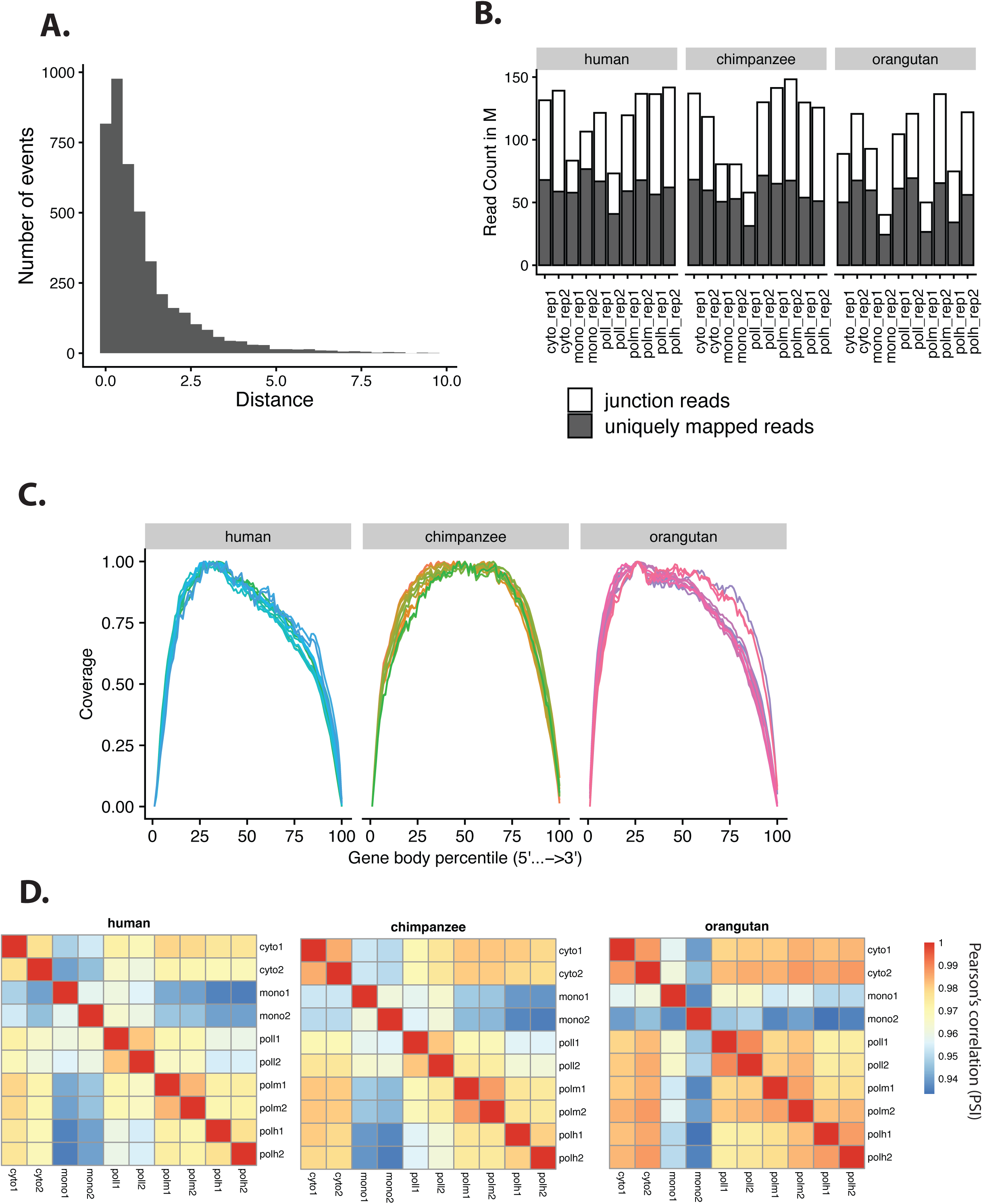
A) Distribution of cumulative manhattan distance scores across AS-TC events. B) Number of uniquely mapped reads and junction reads mapped to each fraction collected from human, chimpanzee and orangutan iPSCs. C) Gene coverage in human, chimpanzee, and orangutan sequencing data. D) Pearson correlation of PSI values between replicates.

**Supplementary Fig 2.**
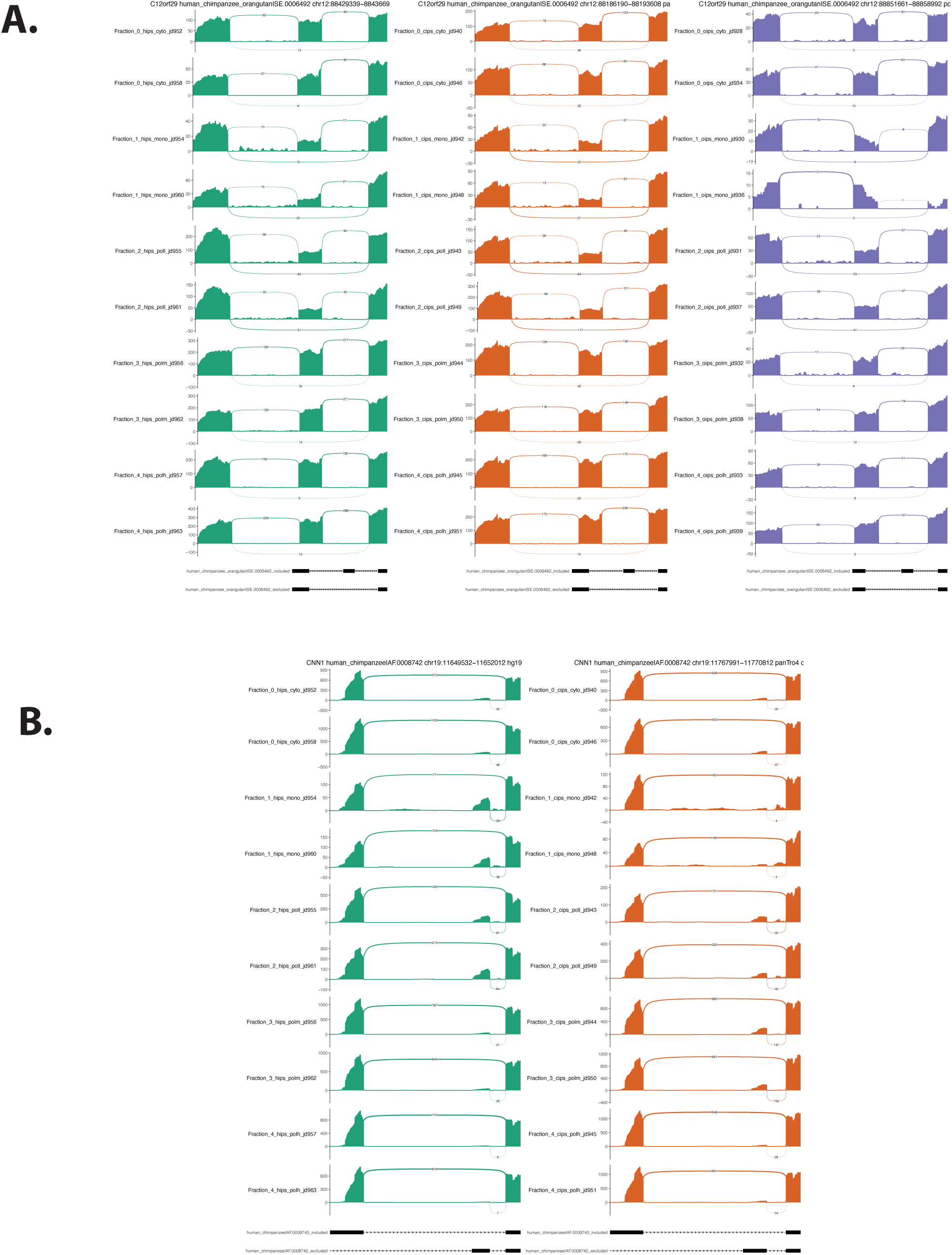
A) Sashimi plot depicting junction reads mapped to a skipped exon event within C12orf29 (as shown in Figure 2E) across all fractions in human (green), chimpanzee (orange) and orangutan (purple). B) Sashimi plot depicting junction reads mapped to an alternative first exon event within CNN1 (as shown in Figure 2F) across all fractions in human (green) and chimpanzee (orange).

**Supplementary Fig 3.**
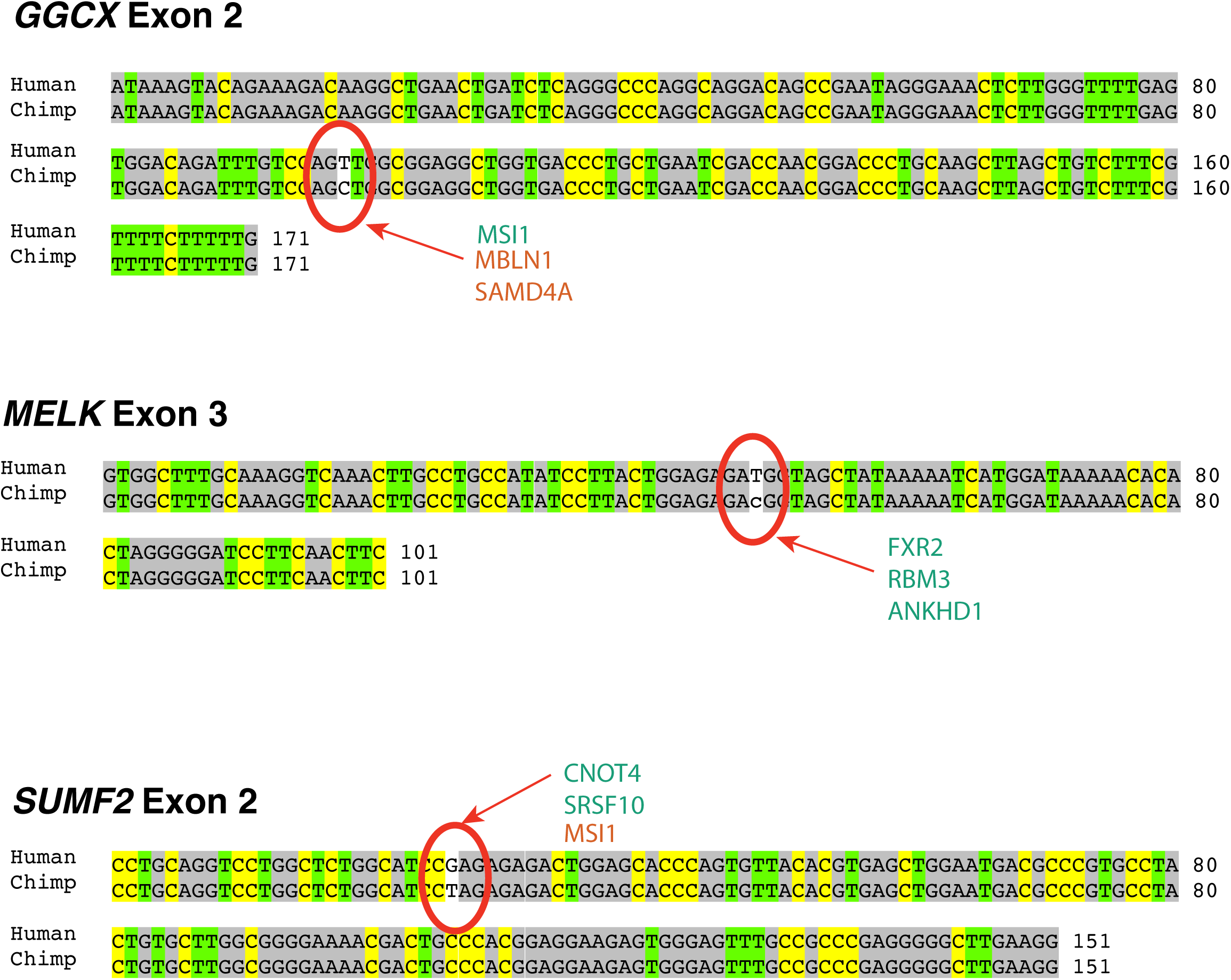
Sequence alignments of orthologous skipped exons from human and chimpanzee frac seq data. (A) Alignment of GGCX exon 2. (B) Alignment of SUMF2 exon 2. (C) Alignment of MELK Exon 3. RBPs predicted to interact with either the human or chimpanzee allele are show in green or orange font, respectively.

**Supplementary Fig 4.**
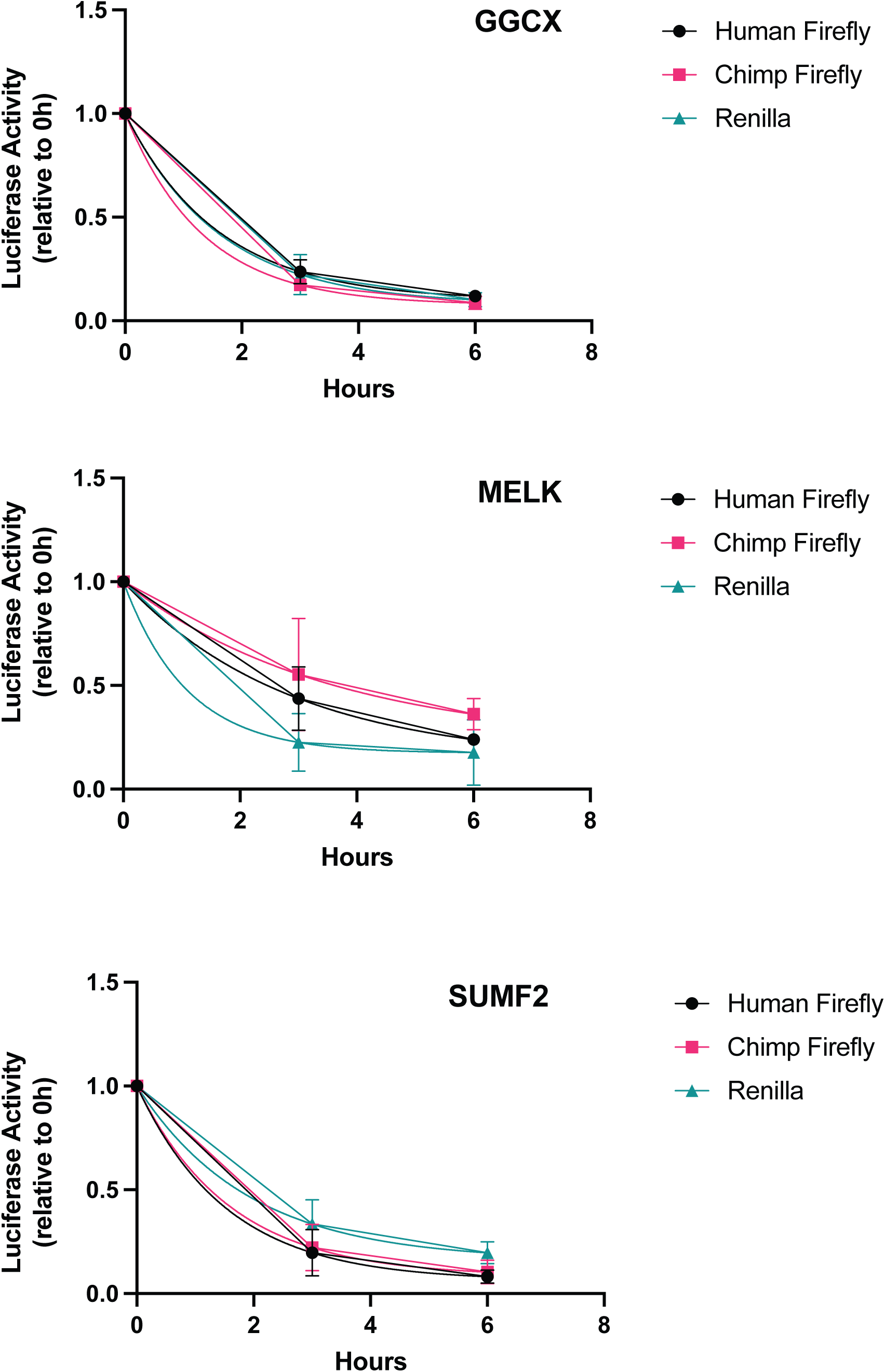
Luciferase enzyme activity stability measurements. HEK293T cells were transfected with human/chimp orthologous firefly reporters and a Renilla luciferase reporter. 48 hours post transfection cells were treated with cycloheximide for 0, 3 or 6 hours. Luciferase levels were measured by dual luciferase assay and normalized to the 0h time point. Data were fit to a one phase decay nonlinear model and decay constants (K) for human and chimp ortholog pairs were compared using an Extra Sum of Squares F-test. The reporter pairs did not exhibit significant differences in decay constants.

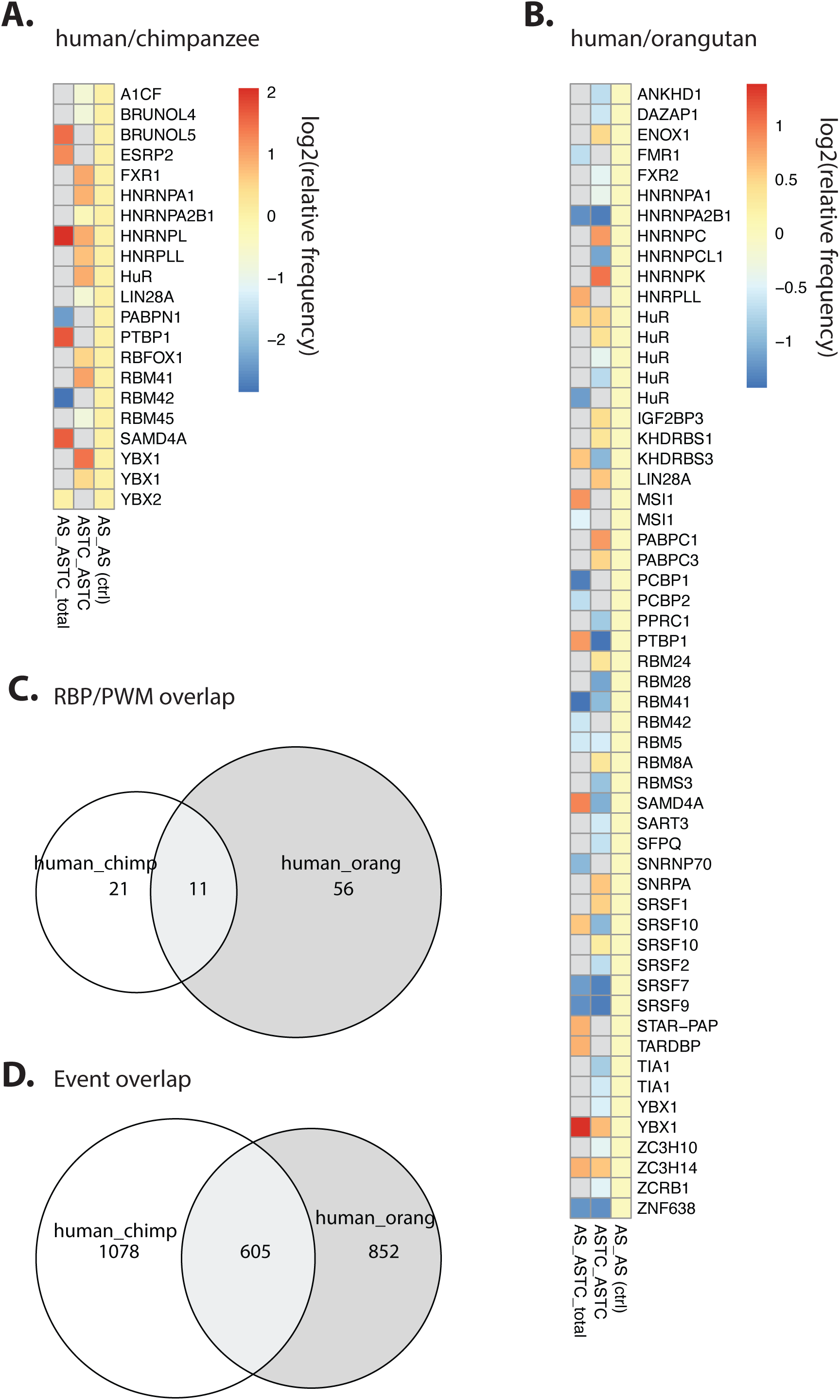

## References

1. Barberan-Soler S, Lambert NJ, Zahler AM. 2009. Global analysis of alternative splicing uncovers developmental regulation of nonsense-mediated decay in C. elegans. RNA 15: 1652–1660.

2. Barbosa-Morais NL, Irimia M, Pan Q, Xiong HY, Gueroussov S, Lee LJ, Slobodeniuc V, Kutter C, Watt S, Colak R, et al. 2012. The evolutionary landscape of alternative splicing in vertebrate species. Science 338: 1587–1593.

3. Blair JD, Hockemeyer D, Doudna JA, Bateup HS, Floor SN. 2017. Widespread Translational Remodeling during Human Neuronal Differentiation. Cell Rep 21: 2005–2016.

4. Blaustein M, Pelisch F, Tanos T, Muñoz MJ, Wengier D, Quadrana L, Sanford JR, Muschietti JP, Kornblihtt AR, Cáceres JF, et al. 2005. Concerted regulation of nuclear and cytoplasmic activities of SR proteins by AKT. Nat Struct Mol Biol 12: 1037–1044.

5. Bustin SA, Benes V, Garson JA, Hellemans J, Huggett J, Kubista M, Mueller R, Nolan T, Pfaffl MW, Shipley GL, et al. 2009. The MIQE guidelines: minimum information for publication of quantitative real-time PCR experiments. Clin Chem 55: 611–622.

6. Calarco JA, Xing Y, Cáceres M, Calarco JP, Xiao X, Pan Q, Lee C, Preuss TM, Blencowe BJ. 2007. Global analysis of alternative splicing differences between humans and chimpanzees. Genes Dev 21: 2963–2975.

7. Chemla Y, Peeri M, Heltberg ML, Eichler J, Jensen MH, Tuller T, Alfonta L. 2020. A possible universal role for mRNA secondary structure in bacterial translation revealed using a synthetic operon. Nat Commun 11: 4827.

8. Dobin A, Davis CA, Schlesinger F, Drenkow J, Zaleski C, Jha S, Batut P, Chaisson M, Gingeras TR. 2013. STAR: ultrafast universal RNA-seq aligner. Bioinformatics 29: 15–21.

9. ENCODE Project Consortium. 2012. An integrated encyclopedia of DNA elements in the human genome. Nature 489: 57–74.

10. Fiddes IT, Armstrong J, Diekhans M, Nachtweide S, Kronenberg ZN, Underwood JG, Gordon D, Earl D, Keane T, Eichler EE, et al. 2018. Comparative Annotation Toolkit (CAT)— simultaneous clade and personal genome annotation. Genome Res 28: 1029–1038.

11. Field AR, Jacobs FMJ, Fiddes IT, Phillips APR, Reyes-Ortiz AM, LaMontagne E, Whitehead L, Meng V, Rosenkrantz JL, Haeussler M, et al. 2017. Structurally conserved primate lncRNAs are transiently expressed during human cortical differentiation and influence cell type specific genes. http://dx.doi.org/10.1101/232553.

12. Floor SN, Doudna JA. 2016a. Tunable protein synthesis by transcript isoforms in human cells. eLife 5. http://dx.doi.org/10.7554/elife.10921.

13. Floor SN, Doudna JA. 2016b. Tunable protein synthesis by transcript isoforms in human cells. Elife 5. http://dx.doi.org/10.7554/eLife.10921.

14. Fu X-D, Ares M Jr. 2014. Context-dependent control of alternative splicing by RNA-binding proteins. Nat Rev Genet 15: 689.

15. Gallego-Paez LM, Bordone MC, Leote AC, Saraiva-Agostinho N, Ascensão-Ferreira M, Barbosa-Morais NL. 2017. Alternative splicing: the pledge, the turn, and the prestige : The key role of alternative splicing in human biological systems. Hum Genet 136: 1015–1042.

16. Gebauer F, Preiss T, Hentze MW. 2012. From cis-regulatory elements to complex RNPs and back. Cold Spring Harb Perspect Biol 4: a012245.

17. Gu W, Zhou T, Wilke CO. 2010. A universal trend of reduced mRNA stability near the translation-initiation site in prokaryotes and eukaryotes. PLoS Comput Biol 6: e1000664.

18. Huang Y, Steitz JA. 2001. Splicing factors SRp20 and 9G8 promote the nucleocytoplasmic export of mRNA. Mol Cell 7: 899–905.

19. Kent WJ, Sugnet CW, Furey TS, Roskin KM, Pringle TH, Zahler AM, Haussler a. D. 2002. The Human Genome Browser at UCSC. Genome Research 12: 996–1006. http://dx.doi.org/10.1101/gr.229102.

20. Khan Z, Ford MJ, Cusanovich DA, Mitrano A, Pritchard JK, Gilad Y. 2013. Primate transcript and protein expression levels evolve under compensatory selection pressures. Science 342: 1100–1104.

21. Langmead B, Trapnell C, Pop M, Salzberg SL. 2009. Ultrafast and memory-efficient alignment of short DNA sequences to the human genome. Genome Biol 10: R25.

22. Lareau LF, Brenner SE. 2015. Regulation of splicing factors by alternative splicing and NMD is conserved between kingdoms yet evolutionarily flexible. Mol Biol Evol 32: 1072–1079.

23. Lareau LF, Inada M, Green RE, Wengrod JC, Brenner SE. 2007. Unproductive splicing of SR genes associated with highly conserved and ultraconserved DNA elements. Nature 446: 926–929.

24. Leclair NK, Brugiolo M, Urbanski L, Lawson SC, Thakar K, Yurieva M, George J, Hinson JT, Cheng A, Graveley BR, et al. 2020. Poison Exon Splicing Regulates a Coordinated Network of SR Protein Expression during Differentiation and Tumorigenesis. Mol Cell 80: 648–665.e9.

25. Le Hir H, Gatfield D, Izaurralde E, Moore MJ. 2001. The exon-exon junction complex provides a binding platform for factors involved in mRNA export and nonsense-mediated mRNA decay. EMBO J 20: 4987–4997.

26. Lewis BP, Green RE, Brenner SE. 2003. Evidence for the widespread coupling of alternative splicing and nonsense-mediated mRNA decay in humans. Proc Natl Acad Sci U S A 100: 189–192.

27. Lu S, Cullen BR. 2003. Analysis of the stimulatory effect of splicing on mRNA production and utilization in mammalian cells. RNA 9: 618–630.

28. Mao Y, Liu H, Liu Y, Tao S. 2014. Deciphering the rules by which dynamics of mRNA secondary structure affect translation efficiency in Saccharomyces cerevisiae. Nucleic Acids Res 42: 4813–4822.

29. Maslon MM, Heras SR, Bellora N, Eyras E, Cáceres JF. 2014. The translational landscape of the splicing factor SRSF1 and its role in mitosis. Elife e02028.

30. Mauger DM, Cabral BJ, Presnyak V, Su SV, Reid DW, Goodman B, Link K, Khatwani N, Reynders J, Moore MJ, et al. 2019. mRNA structure regulates protein expression through changes in functional half-life. Proc Natl Acad Sci U S A 116: 24075–24083.

31. Mazin PV, Jiang X, Fu N, Han D, Guo M, Gelfand MS, Khaitovich P. 2018. Conservation, evolution, and regulation of splicing during prefrontal cortex development in humans, chimpanzees, and macaques. RNA 24: 585–596.

32. Merkin J, Russell C, Chen P, Burge CB. 2012. Evolutionary dynamics of gene and isoform regulation in Mammalian tissues. Science 338: 1593–1599.

33. Michlewski G, Sanford JR, Cáceres JF. 2008. The splicing factor SF2/ASF regulates translation initiation by enhancing phosphorylation of 4E-BP1. Mol Cell 30: 179–189.

34. Müller-McNicoll M, Botti V, de Jesus Domingues AM, Brandl H, Schwich OD, Steiner MC, Curk T, Poser I, Zarnack K, Neugebauer KM. 2016. SR proteins are NXF1 adaptors that link alternative RNA processing to mRNA export. Genes Dev 30: 553–566.

35. Niederer RO, Rojas-Duran MF, Zinshteyn B, Gilbert WV. 2022. Direct analysis of ribosome targeting illuminates thousand-fold regulation of translation initiation. Cell Syst 13: 256–264.e3.

36. Ni JZ, Grate L, Donohue JP, Preston C, Nobida N, O’Brien G, Shiue L, Clark TA, Blume JE, Ares M Jr. 2007. Ultraconserved elements are associated with homeostatic control of splicing regulators by alternative splicing and nonsense-mediated decay. Genes Dev 21: 708–718.

37. Nott A, Le Hir H, Moore MJ. 2004. Splicing enhances translation in mammalian cells: an additional function of the exon junction complex. Genes Dev 18: 210–222.

38. Pagès H, Aboyoun P, Gentleman R, DebRoy S. 2020. Biostrings: Efficient manipulation of biological strings. R package version 2.48. 0.

39. Pan Q, Saltzman AL, Kim YK, Misquitta C, Shai O, Maquat LE, Frey BJ, Blencowe BJ. 2006. Quantitative microarray profiling provides evidence against widespread coupling of alternative splicing with nonsense-mediated mRNA decay to control gene expression. Genes Dev 20: 153–158.

40. Pertea M, Pertea GM, Antonescu CM, Chang T-C, Mendell JT, Salzberg SL. 2015. StringTie enables improved reconstruction of a transcriptome from RNA-seq reads. Nat Biotechnol 33: 290–295.

41. Pfeiffer BD, Truman JW, Rubin GM. 2012. Using translational enhancers to increase transgene expression in Drosophila. Proc Natl Acad Sci U S A 109: 6626–6631.

42. Presnyak V, Alhusaini N, Chen Y-H, Martin S, Morris N, Kline N, Olson S, Weinberg D, Baker KE, Graveley BR, et al. 2015. Codon optimality is a major determinant of mRNA stability. Cell 160: 1111–1124.

43. Ray D, Ha KCH, Nie K, Zheng H, Hughes TR, Morris QD. 2017. RNAcompete methodology and application to determine sequence preferences of unconventional RNA-binding proteins. Methods 118-119: 3–15.

44. Ray D, Kazan H, Chan ET, Peña Castillo L, Chaudhry S, Talukder S, Blencowe BJ, Morris Q, Hughes TR. 2009. Rapid and systematic analysis of the RNA recognition specificities of RNA-binding proteins. Nat Biotechnol 27: 667–670.

45. Ray D, Kazan H, Cook KB, Weirauch MT, Najafabadi HS, Li X, Gueroussov S, Albu M, Zheng H, Yang A, et al. 2013. A compendium of RNA-binding motifs for decoding gene regulation. Nature 499: 172–177.

46. Reixachs-Solé M, Ruiz-Orera J, Albà MM, Eyras E. 2020. Ribosome profiling at isoform level reveals evolutionary conserved impacts of differential splicing on the proteome. Nat Commun 11: 1768.

47. Saltzman AL, Kim YK, Pan Q, Fagnani MM, Maquat LE, Blencowe BJ. 2008. Regulation of multiple core spliceosomal proteins by alternative splicing-coupled nonsense-mediated mRNA decay. Mol Cell Biol 28: 4320–4330.

48. Sanford JR. 2004. A novel role for shuttling SR proteins in mRNA translation. Genes & Development 18: 755–768. http://dx.doi.org/10.1101/gad.286404.

49. Sanford JR, Gray NK, Beckmann K, Cáceres JF. 2004. A novel role for shuttling SR proteins in mRNA translation. Genes Dev 18: 755–768.

50. Siepel A. 2005. Evolutionarily conserved elements in vertebrate, insect, worm, and yeast genomes. Genome Research 15: 1034–1050. http://dx.doi.org/10.1101/gr.3715005.

51. Siepel A, Haussler D. Phylogenetic Hidden Markov Models. Statistical Methods in Molecular Evolution 325–351. http://dx.doi.org/10.1007/0-387-27733-1_12.

52. Sterne-Weiler T, Martinez-Nunez RT, Howard JM, Cvitovik I, Katzman S, Tariq MA, Pourmand N, Sanford JR. 2013. Frac-seq reveals isoform-specific recruitment to polyribosomes. Genome Res 23: 1615–1623.

53. Swartz JE, Bor Y-C, Misawa Y, Rekosh D, Hammarskjold M-L. 2007. The Shuttling SR Protein 9G8 Plays a Role in Translation of Unspliced mRNA Containing a Constitutive Transport Element. Journal of Biological Chemistry 282: 19844–19853. http://dx.doi.org/10.1074/jbc.m701660200.

54. Taliaferro JM, Vidaki M, Oliveira R, Olson S, Zhan L, Saxena T, Wang ET, Graveley BR, Gertler FB, Swanson MS, et al. 2016. Distal Alternative Last Exons Localize mRNAs to Neural Projections. Mol Cell 61: 821–833.

55. Thomas JD, Polaski JT, Feng Q, De Neef EJ, Hoppe ER, McSharry MV, Pangallo J, Gabel AM, Belleville AE, Watson J, et al. 2020. RNA isoform screens uncover the essentiality and tumor-suppressor activity of ultraconserved poison exons. Nat Genet 52: 84–94.

56. Tuller T, Carmi A, Vestsigian K, Navon S, Dorfan Y, Zaborske J, Pan T, Dahan O, Furman I, Pilpel Y. 2010. An evolutionarily conserved mechanism for controlling the efficiency of protein translation. Cell 141: 344–354.

57. Wang ET, Cody NAL, Jog S, Biancolella M, Wang TT, Treacy DJ, Luo S, Schroth GP, Housman DE, Reddy S, et al. 2012. Transcriptome-wide regulation of pre-mRNA splicing and mRNA localization by muscleblind proteins. Cell 150: 710–724.

58. Wasserman WW, Sandelin A. 2004. Applied bioinformatics for the identification of regulatory elements. Nature Reviews Genetics 5: 276–287. http://dx.doi.org/10.1038/nrg1315.

59. Wiegand HL, Lu S, Cullen BR. 2003. Exon junction complexes mediate the enhancing effect of splicing on mRNA expression. Proc Natl Acad Sci U S A 100: 11327–11332.

60. Wong QW-L, Vaz C, Lee QY, Zhao TY, Luo R, Archer SK, Preiss T, Tanavde V, Vardy LA. 2016. Embryonic Stem Cells Exhibit mRNA Isoform Specific Translational Regulation. PLoS One 11: e0143235.

61. Woodward LA, Mabin JW, Gangras P, Singh G. 2017. The exon junction complex: a lifelong guardian of mRNA fate. Wiley Interdiscip Rev RNA 8. http://dx.doi.org/10.1002/wrna.1411

